# 25-hydroxycholesterol mediates brain cytokine production and neutrophil infiltration in a mouse model of lipopolysaccharide-induced neuroinflammation

**DOI:** 10.1101/2023.10.09.561598

**Authors:** Johnathan Romero, Danira Toral-Rios, Jinsheng Yu, Steven M Paul, Anil G Cashikar

## Abstract

Neuroinflammation has been implicated in the pathogenesis of several neurologic and psychiatric disorders. Microglia are key drivers of neuroinflammation and in response to different inflammatory stimuli overexpress a proinflammatory signature of genes. Among these, *Ch25h* is a gene overexpressed in brain tissue from Alzheimer’s disease as well as various mouse models of neuroinflammation. *Ch25h* encodes cholesterol 25-hydroxylase, an enzyme that hydroxylates cholesterol to form 25-hydroxycholesterol (25HC). 25HC, an immune-oxysterol primarily produced by activated microglia, is further metabolized to 7α,25-dihydroxycholesterol, which is a potent chemoattractant for leukocytes. We have also previously shown that 25HC increases production and secretion of the proinflammatory cytokine, IL-1β, by mouse microglia treated with lipopolysaccharide (LPS). In the present study, wildtype (*WT*) and *Ch25h*-knockout (*CKO*) mice were peripherally administered LPS to induce an inflammatory state in the brain. In LPS-treated *WT* mice, *Ch25h* expression and 25HC levels increased in brain relative to vehicle-treated *WT* mice. Interestingly, 25HC levels were significantly higher in LPS-treated *WT* female compared to male mice. Activation of microglia and astrocytes in response to systemic LPS was suppressed in *CKO* mice relative to *WT* mice. Proinflammatory cytokine production and intra-parenchymal infiltration of neutrophils strongly correlated with brain 25HC levels and were significantly lower in *CKO* compared to *WT* mice. Overall, our results show that 25HC mediates a sex-specific proinflammatory response in the brain characterized by activation of both microglia and astrocytes but also by neutrophil migration into the brain parenchyma presumably due to 7α,25-diHC produced from 25HC.

## BACKGROUND

Emerging evidence indicates that neuroinflammation plays an important role in many neurological and psychiatric disorders^1,2^. Therefore, elucidating the cellular pathways and chemical mediators involved in neuroinflammation could shed light on promising therapeutic targets. Changes in CNS lipid metabolism, including the synthesis of various cholesterol metabolites, such as bile acids, steroid hormones, and oxysterols, have all been shown to influence immune regulation and neuroinflammation^3^. One oxysterol in particular, 25-hydroxycholesterol (25HC), has especially potent pro- and anti-inflammatory properties under various conditions^4,5^, as well as anti-viral properties against a variety of viruses^6^. 25HC is produced via the enzymatic hydroxylation of cholesterol by cholesterol 25-hydroxylase (CH25H). The *Ch25h* gene is primarily expressed in myeloid cells when challenged with inflammatory stimuli. In the central nervous system (CNS), *Ch25h* is expressed mainly by microglia in mice and humans^7,8^ under inflammatory conditions^9^. Importantly, *Ch25h* is also among a set of genes expressed by disease-associated microglia (DAM) that are chronically upregulated in various neurodegenerative diseases^10,11^. Taken together, these findings suggest that the principal function of CH25H in the CNS relates to innate immunity and immune surveillance. We and others have postulated that prolonged expression of the enzyme and increased 25HC production may contribute to neuroinflammation and neurodegeneration^12^.

The cellular mechanisms by which 25HC regulates immune function appear to stem from its ability to strongly influence cholesterol metabolism, including suppression of cholesterol biosynthesis and enhancement of cholesterol esterification^13,14^. Recent studies have provided conflicting evidence about the function of CH25H in modulating inflammation, with some studies reporting greater pro-inflammatory cytokine release^5^ and others finding less pro-inflammatory cytokine expression in *Ch25h*-deficient mice^12,16,17^. In murine bone-marrow derived macrophages, the SREBP2-SCAP complex, an inhibitory target of 25HC involved in cholesterol metabolism, was found to facilitate inflammasome activation, suggesting an anti-inflammatory role for CH25H^18^. However, in murine brain-resident macrophages (i.e., microglia), 25HC increased interleukin-1-beta (IL-1β) secretion, indicating a potential pro-inflammatory role for CH25H in the CNS^12^. Known potent inducers of *Ch25h* expression include ligands of type I interferon-receptors (IFNRs) and Toll-like receptors (TLRs), particularly TLR4^19^.

Lipopolysaccharide (LPS), a surface glycolipid found in most gram-negative bacteria, is a TLR4 agonist that not only increases *Ch25h* expression^20^, but also triggers many of the inflammatory processes involved in neurodegeneration, including cytokine release, gliosis, and peripheral leukocyte infiltration. Recently, neuroinflammation has been extensively studied in the context of mouse models of neurodegeneration and demyelination^21,22^. Neuroinflammation may also be acutely induced by peripheral injection of LPS that mimics bacterial infection^23^.

To better understand the role of CH25H and 25HC in neuroinflammation, we administered LPS peripherally to induce neuroinflammation in *Ch25h* knockout and wildtype female and male mice. We observed a striking correlation of brain 25HC and cytokine levels, as well as with the infiltration of neutrophils into the CNS. Our results also suggest that brain 25HC levels may underlie the greater levels of LPS-induced neuroinflammation observed in female relative to male mice.

## RESULTS

### Intraperitoneal administration of LPS induces *Ch25h* expression and 25HC synthesis in the brain

To understand how CH25H and 25HC influences neuroinflammation, we intraperitoneally (i.p) administered LPS or vehicle (PBS) twice (two-hit LPS model) in wildtype (*WT*) and *ch25h-/-* (*CKO*) mice (Fig 1A). This LPS-mediated neuroinflammation model was recently shown to result in increased cytokine levels as well as neutrophil infiltration in the brain^24^. Expression of *Ch25h* was confirmed in the *WT* and *CKO* by qPCR. Replicating findings from previous studies^20,25,26^, LPS administration promoted an upregulation of *Ch25h* expression in *WT* mice by almost 10-fold (p<0.0001) and, as expected, no expression in the *CKO* group was observed (Fig. 1B). Based on single cell transcriptomics studies^7,8,27^ it is likely that *Ch25h* overexpression is mainly due to activated microglia (also see [The Myeloid Landscape 2](http://research-pub.gene.com/BrainMyeloidLandscape/BrainMyeloidLandscape2/#Mouse-gene/Mouse-gene/12642/geneReport.html)).

**Figure 1:**
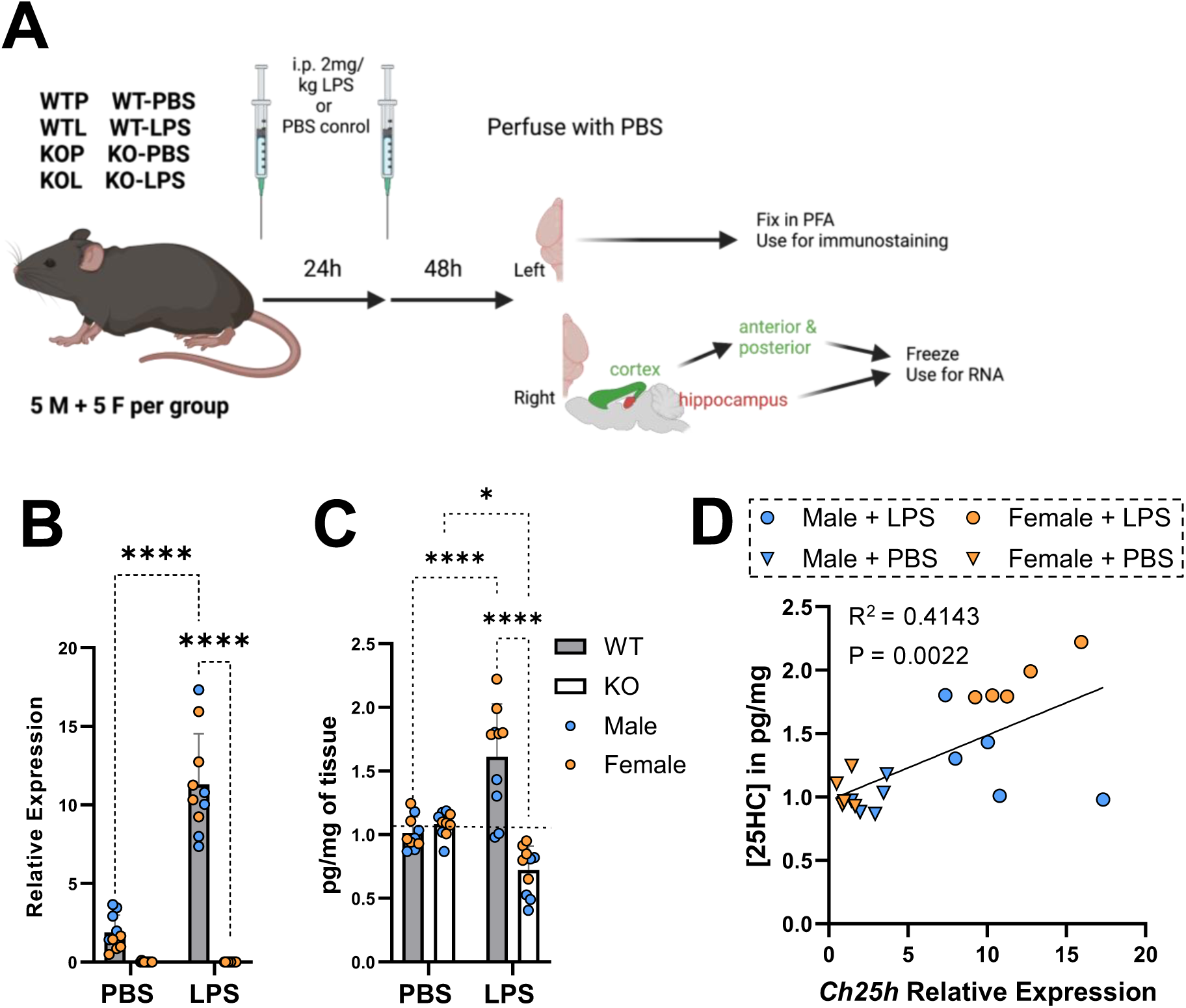
Mouse model for LPS-mediated neuroinflammation and quantification of 25HC and *Ch25h* expression. (A) Schematic of 2-hit LPS mouse model. (B) Relative expression of *Ch25h* and (C) measured concentrations of 25HC in PBS- and LPS-injected wildtype (WT) and Ch25h- deficient (KO) mouse brains. Male mice are indicated in blue and female mice are in orange. Data are presented as mean values ± SD. Statistical significance was defined using a two-way ANOVA with Tukey’s post-hoc test (\**P* < 0.05, \*\*\**P* < 0.001, & \*\*\*\**P* < 0.0001). (D) Correlation plot of Ch25h mRNA expression and 25HC produced in the brain tissue. Male mice are indicated in blue and female mice are in orange; PBS controls are inverted triangles and LPS- treated mice are circles – as shown at the top of the correlation plot. All points were fit by linear regression and the R^2^ and P values are shown at the tope left.

To confirm that the increase in *Ch25h* expression resulted in an increase in 25HC synthesis, the relative concentrations of 25HC (as a ratio of the total internal standard of 25HC) was quantified in the cortical tissue by liquid chromatography/mass spectrometry (LC/MS) (Fig. 1C). Consistent with expression of *Ch25h*, a significant increase (p<0.0001) in the relative amounts of 25HC was observed in the LPS-treated *WT* compared to the PBS-treated *WT* mice, while the *CKO* cohort demonstrated levels of 25HC that were lower or comparable to the PBS control. The low levels of 25HC observed in brain tissue from PBS-treated *WT* mice and in *CKO* mice are presumably produced by pathways independent of CH25H enzyme activity such as cholesterol hydroxylation by CYP3A4 as reported earlier^28^. Increased production of 25HC in LPS-treated samples correlates highly with *Ch25h* mRNA expression (Fig 1D) indicating that the 25HC synthesized after LPS-treatment resulted from Ch25h enzyme activity. To ensure that changes in structurally isomeric oxysterols did not skew the 25HC measurements, we also determined the concentrations of the two isomeric oxysterols – namely, 24S-hydroxycholesterol (24(S)HC) and 27-hydroxycholesterol (27HC) – that are appreciably present in the CNS (Figures S1A & B). Confirming that LPS selectively induces 25HC synthesis, treatment with LPS did not increase levels of 24(S)HC or 27HC. Thus, we conclude that peripheral administration of LPS induces the expression of *Ch25h* in the brain, leading to elevated 25HC synthesis.

Interestingly, LPS-induced 25HC synthesis was significantly higher in female than in male mice. Since no significant difference was obvious between males and females in the expression of *Ch25h* mRNA, we tested whether 7α-hydroxylation of 25HC accounts for the difference between males and females by examining the expression of *Cyp7b1*, the gene encoding the enzyme for 7α-hydroxylation of 25HC to produce 7α,25-dihydroxycholesterol. No significant dfifferences in *Cyp7b1* expression were apparent (Fig S2). Our results suggest that 25HC may be metabolized differently in males and females leading to differences in 25HC levels after LPS challenge.

### Ch25h deficiency reduces gliosis induced by LPS administration

Activation of brain-resident immune cells (i.e., microglia) is associated with neuroinflammation. To assess the role of CH25H in promoting microglial activation, hippocampal sections of wildtype and *Ch25h*-deficient mice were probed for markers of microglial activation by immunostaining for ionized calcium-binding adaptor molecule 1 (IBA1) as well as the microglial phagocytosis marker cluster of differentiation 68 (CD68), expressed by activated microglia (Figures 2A & B). As expected, LPS treatment significantly increased the area covered by IBA1 (p<0.001) and CD68 (p<0.0001) immunoreactivity in *WT* mice compared with the PBS-treated *WT* mice, while no significant effect of *Ch25h* genotype on IBA1 immunoreactivity was observed (Figure 2A & B). However, CD68 immunoreactivity was substantially lower in *Ch25h*-deficient mouse hippocampi, suggesting that microglial activation is partially elevated by *Ch25h* (Fig 2C & D). Additionally, no sex differences in IBA1 or CD68 immunoreactivity were observed.

**Figure 2:**
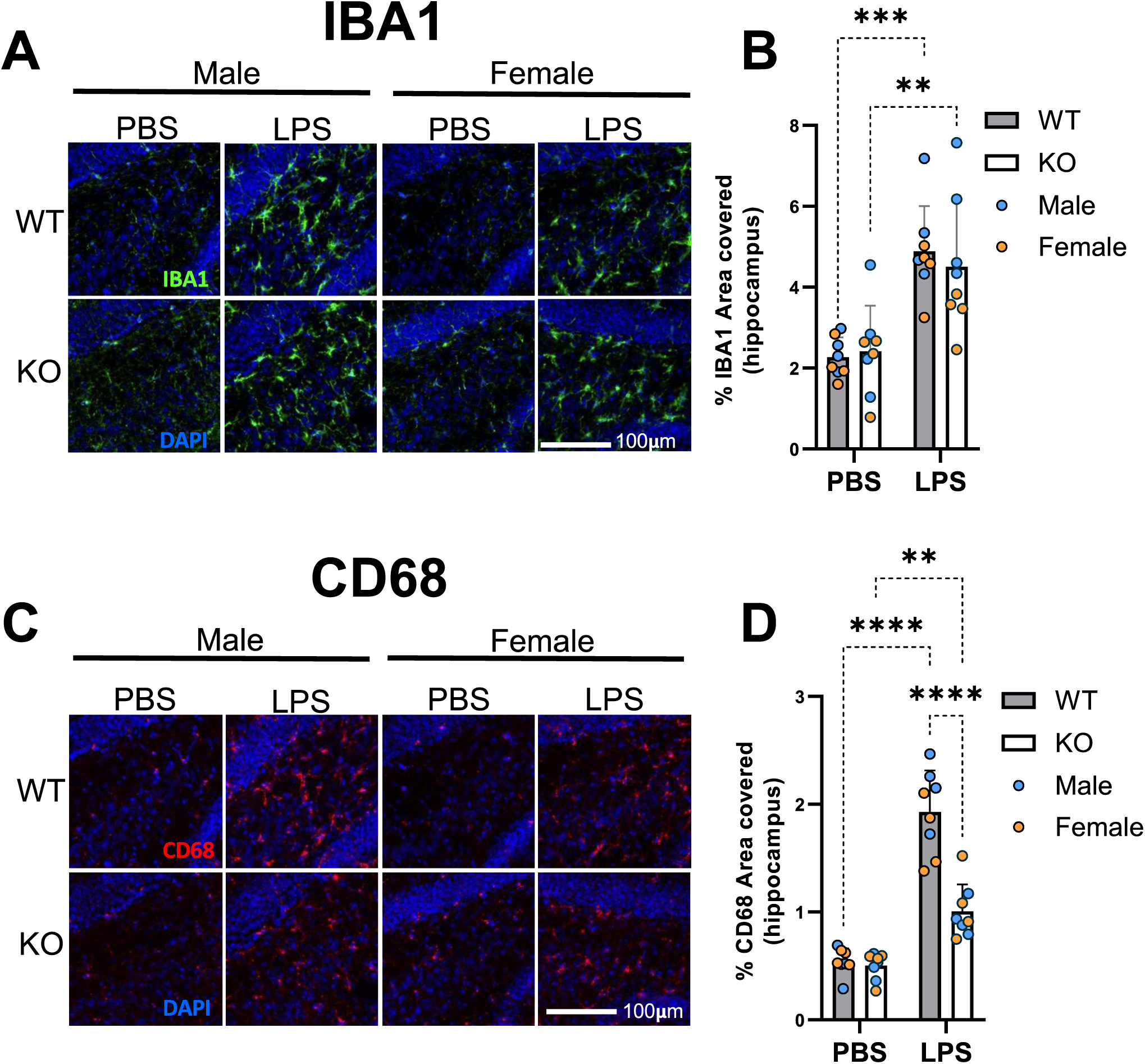
Microglial activation triggered by LPS treatment was mitigated by loss of Ch25h. Representative images of immunostaining (A) and quantification (B) by percentage of IBA1 area in the dentate gyrus. Representative images of immunostaining (C) and quantification (D) by percentage of CD68 area in the dentate gyrus. Data are presented as mean values ± SD. Statistical significance was defined using a two-way ANOVA with Tukey’s post-hoc test (**P < 0.01, ***P < 0.001, & ****P < 0.0001).

To investigate additional markers of microglial activation, we quantified the relative expression of markers of disease associated microglia (DAM) (Fig 3) – namely, *Trem2* (triggering receptor expressed on myeloid cells 2), *Clec7a* (C-type lectin domain family member 7A), and *Axl* (tyrosine-protein kinase receptor UFO). While no LPS-dependent change in *Trem2* expression was observed in the *WT* or *CKO* samples, expression of *Trem2* mRNA was significantly lower (p<0.05) in the *CKO* samples (Fig. 3A). In LPS-treated *WT* samples, males had lower levels of *Trem2* than females. With LPS treatment, *Clec7a* expression was significantly higher (p<0.05) only in *WT* mice compared to PBS-control mice (Fig. 3B). LPS-dependent upregulation (p <0.001) of *Axl* mRNA was also observed in the *WT* mice but not in *CKO* mice (Fig. 3C). These results suggest that despite similar IBA1 immunoreactivity between *WT* and *CKO* mice, microglial activation was significantly lower in *Ch25h*-deficient mice.

**Figure 3:**
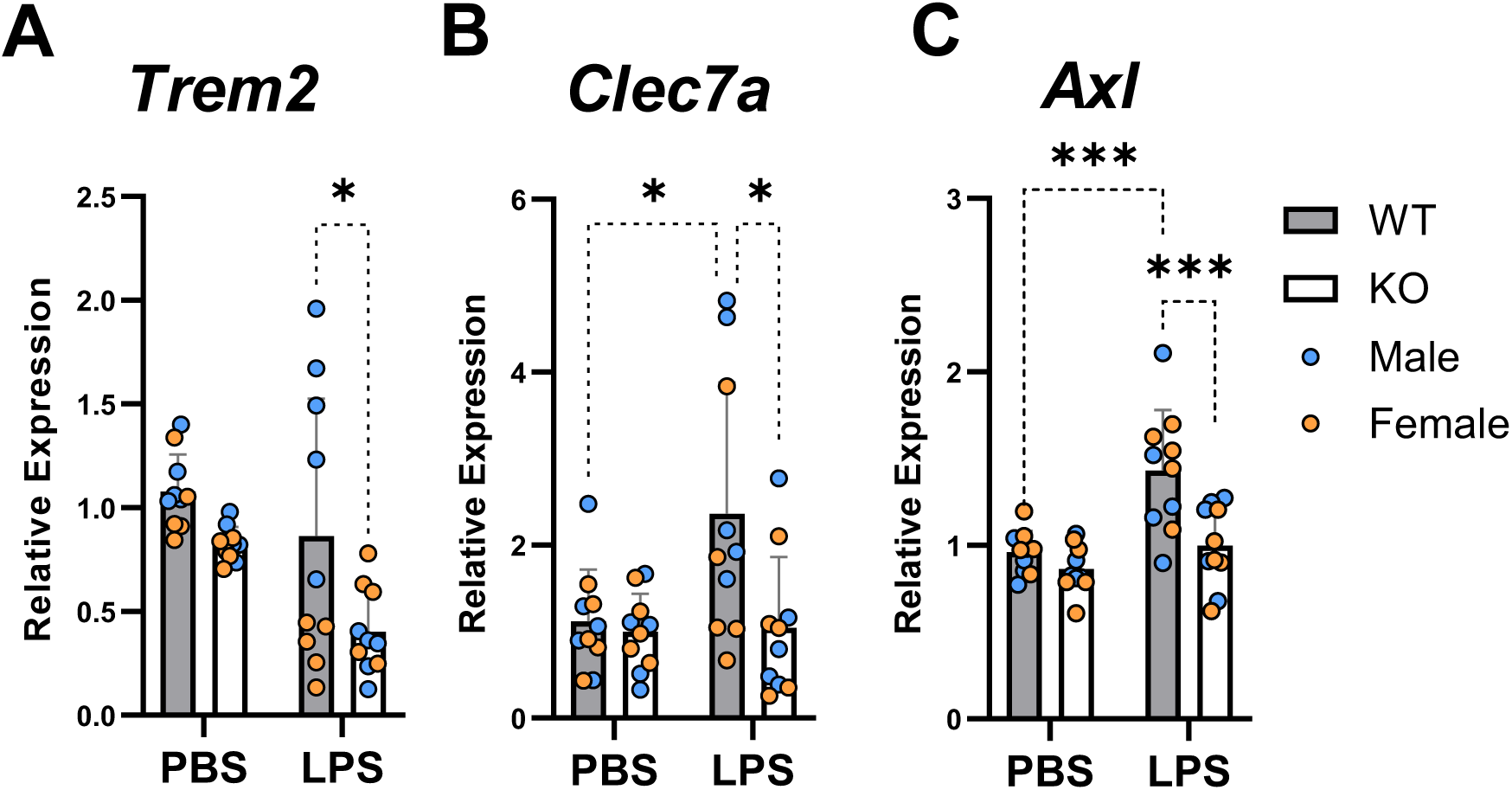
Functional Ch25h promoted gene expression of activated microglia and phagocytic markers upon LPS induction. (A-B) Relative expression of Trem2 (A) Clec7a (B) and Axl (C). Data are presented as mean values ± SD. Statistical significance was defined using a two-way ANOVA with Tukey’s post-hoc test (*P < 0.05, ***P < 0.001).

In addition to microgliosis, serial systemic or local injections of LPS in mice has been shown to induce astrogliosis. Accumulating evidence indicates that altered lipid metabolism is associated with reactive astrogliosis^29,30^. We recently reported that 25HC markedly suppresses cholesterol biosynthesis and increases cholesterol esterification as well as cholesterol efflux via ApoE lipoproteins in murine primary astrocytes^14^ and suggests a possible role for 25HC in astroglial activation. Therefore, to evaluate how 25HC influences astrogliosis *in vivo*, we immunostained mouse brain sections for glial fibrillary acidic marker (GFAP) (Figure 4A). Quantitation of the immunostained images showed a trend (p<0.1686) toward an increase in the percent area of GFAP immunoreactivity in the hippocampi of LPS-treated *WT* mice compared with the PBS-injected mice. Interestingly, among the LPS-injected mice, significantly less area of GFAP immunoreactivity (p<0.05) was observed in the *CKO* cohort (Figure 4B). No differences were detected between *WT* and *CKO* mice treated with PBS. To corroborate these findings on the effect of 25HC in astrogliosis, relative expression of GFAP was quantified in the hippocampus by qPCR. In contrast to the results from the immunofluorescence, relative expression of *Gfap* mRNA increased (p<0.0001) by 2-3-fold in LPS-treated *WT* mice compared to PBS-treated *WT* mice (Figure 4C). Although GFAP expression in LPS-treated *CKO* mice was slightly lower than *WT* mice, the difference was not significant (Figure 4C). Additionally, GFAP levels showed no differences between sexes in protein or mRNA levels.

**Figure 4:**
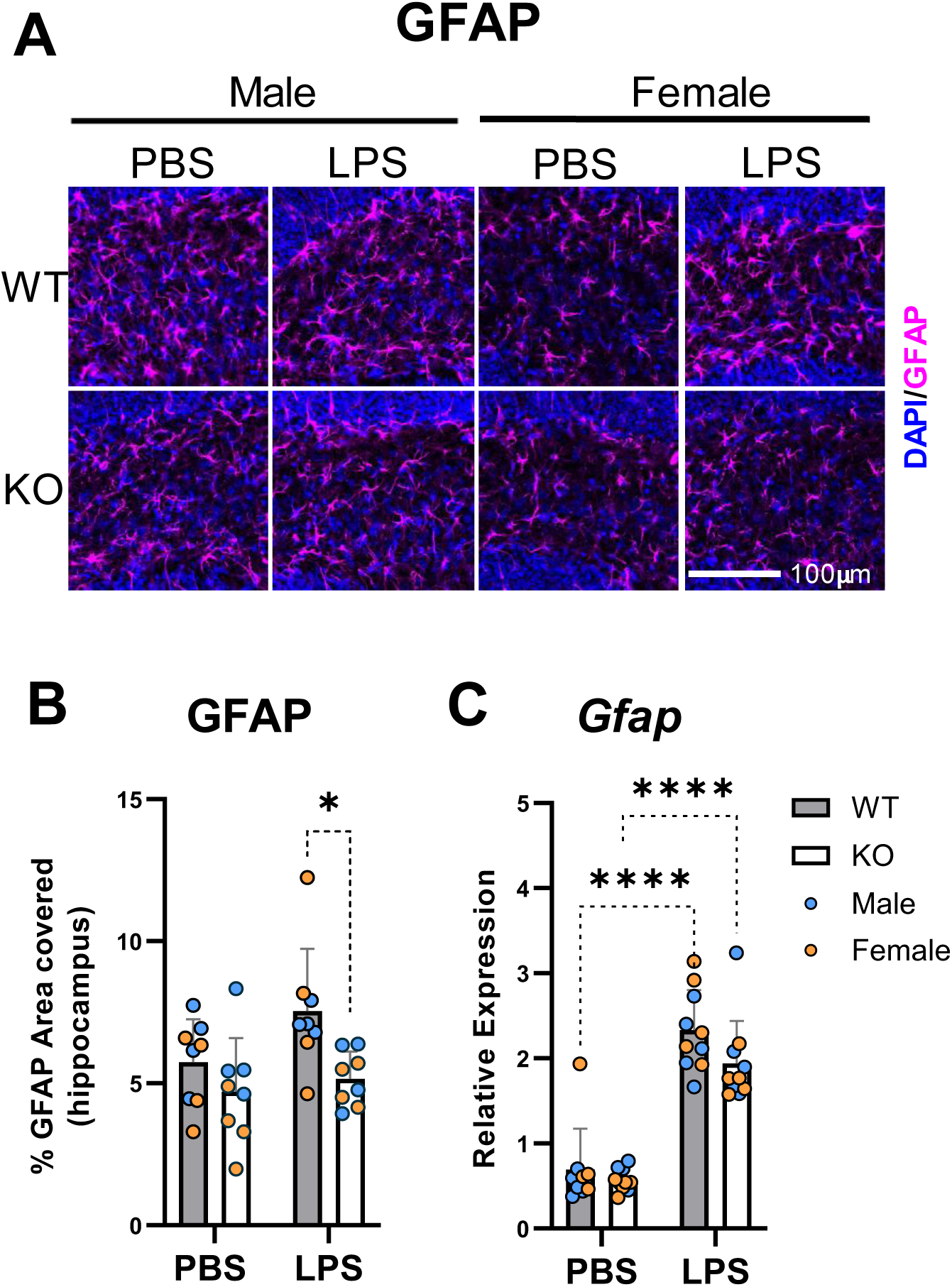
LPS-induced astrogliosis. Representative images of immunostaining (A) and quantification (B) by percentage of GFAP area in the dentate gyrus. C) Relative expression of Gfap mRNA. Data in B and C are presented as mean values ± SD. Statistical significance was defined using a two-way ANOVA with Tukey’s post-hoc test (*P < 0.05, & ****P < 0.0001).

### *Ch25h* deficiency differentially influences LPS-mediated transcriptomic changes in males and females

To understand the underlying mechanisms by which CH25H and 25HC may influence neuroinflammation in this mouse model of LPS-induced neuroinflammation, we carried out transcriptome analysis (RNAseq) of the hippocampal tissue transcriptome. Gene expression in this study is potentially affected by three different variables – namely, treatment (LPS vs PBS), genotype (*CKO* vs *WT*) and sex (females vs males).

Treatment with LPS resulted in major transcriptomic changes in both a genotype- and sex-dependent manner (Supp Table I). While in general LPS treatment resulted in a greater number of differentially expressed genes in females, *Ch25h*-deficiency resulted in fewer gene expression changes in the brains of female mice. When differential gene expression between females and males was examined (Supp table II), female *WT* mice showed greater differential gene expression in response to LPS treatment than male *WT* mice (158 genes up and 66 genes down for fold change >1.5 and q<0.05). Interestingly, such sex-specific differences in the response to LPS were not apparent in *CKO* mice (3 genes up and 4 genes down for fold change >1.5 and q<0.05). The differentially expressed genes in response to LPS treatment are shown as volcano plots in Fig S3. The genes are listed in Supp Table III and the corresponding gene ontology (GO) terms are shown in Fig S4. The differentially upregulated genes following LPS treatment were grouped into GO terms: ‘regulation of defense response’, ‘regulation of cytokine production’, ‘leukocyte migration’ and others (Figure 5).

**Figure 5:**
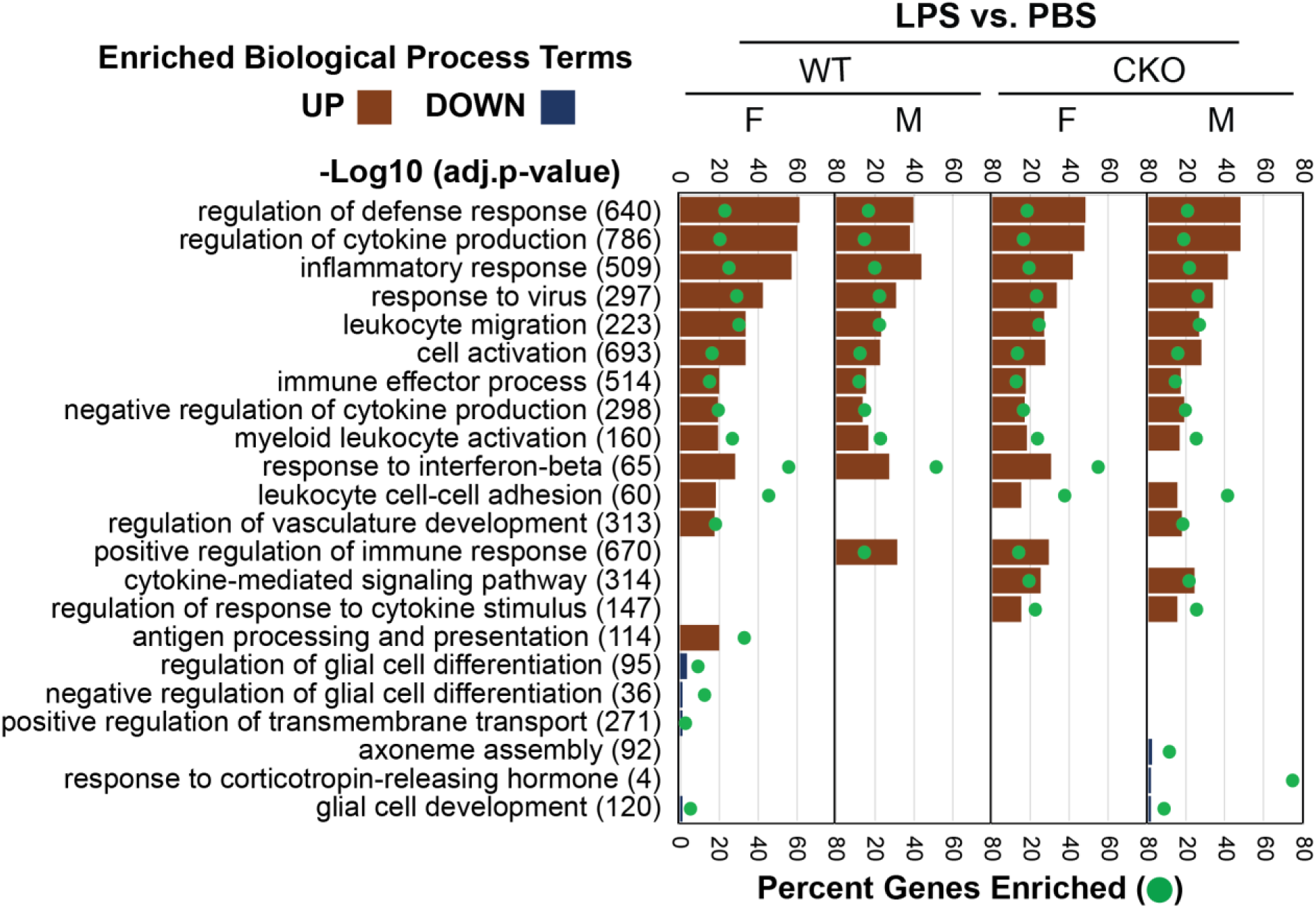
Transcriptomics of mouse hippocampi. Comparison of the LPS effect on female (F) and male (M) mice in the wildtype (WT) and ch25h knockout (CKO) groups. The horizontal brown bars under each group show −log10 (adj. P value) for the gene ontology (GO) biological process on the top X-axis. The green circle shows the percent of genes enriched within each GO term.

To understand whether sex played a role in LPS-mediated neuroinflammation in mice, we compared the LPS response of female and male *WT* and *CKO* mice side-by-side (Figure 5 and Supp Table III). Comparisons of the upregulated pathways in response to LPS treatment (Supp Table IV) revealed that – (i) cytokine production genes included 21.6% hits (log10 q value = − 61.48) in *WT* females; 15.5 % hits (log10 q value = −38.28) in *WT* males; 17.04% hits (log10 q value = −48.16) in *CKO* females; and 19.2% hits (log10 q value = −48.18) in *CKO* males. (ii) leukocyte migration genes included 31.4% hits (log10 q value = −34.45) in *WT* females; 23.31% hits (log10 q value = −23.24) in *WT* males; 25.11% hits (log10 q value = −27.33) in *CKO* females; and 27.35% hits (log10 q value = −26.6) in *CKO* males. These comparisons reveal that LPS treatment triggered significantly greater expression of genes involved in cytokine production and leukocyte migration in *WT* females relative to *WT* males or *CKO* females. Comparisons of the downregulated genes and pathways in response to LPS treatment (Supp Table IV) revealed that regulation of glial differentiation genes included 10.52% hits (log10 q value −3.83) only in *WT* females and no other groups. Furthermore, sex had little impact on the LPS response (up- or down-regulated genes) in *CKO* mice.

Taken together, these results suggest *WT* female mice had a stronger neuroinflammatory response to LPS than *WT* male mice and that these sex-dependent differences in gene expression were absent in *Ch25h*-deficient mice. To further understand this effect, we focused on how Ch25h/25HC influences cytokine production and leukocyte (neutrophil) migration into the brain parenchyma.

### 25HC concentrations correlate with cytokine levels in the cortex

To ascertain the effects of *Ch25h* deficiency on pro-inflammatory cytokine production and gene transcription, we also measured gene expression and secreted cytokine levels for interleukin 1 beta (*Il1b* and IL-1β; Fig 6A, B & C), tumor necrosis factor alpha (*Tnf* and TNFα; Fig 6D, E & F), and interleukin 6 (*Il6* and IL-6; Fig 6G, H & I) by qPCR and ELISA, respectively. LPS treatment resulted in a substantial upregulation of these cytokine mRNAs in both *WT* and *CKO* samples. Expression of *Il1b* increased by nearly 50-fold in LPS-treated *WT* brains relative to PBS-treated controls. Furthermore, *Il1b* expression was significantly lower in LPS-treated *CKO* samples (Fig. 6A), which is consistent with our earlier report with *WT* and *CKO* microglia studied in vitro^12^.

**Figure 6:**
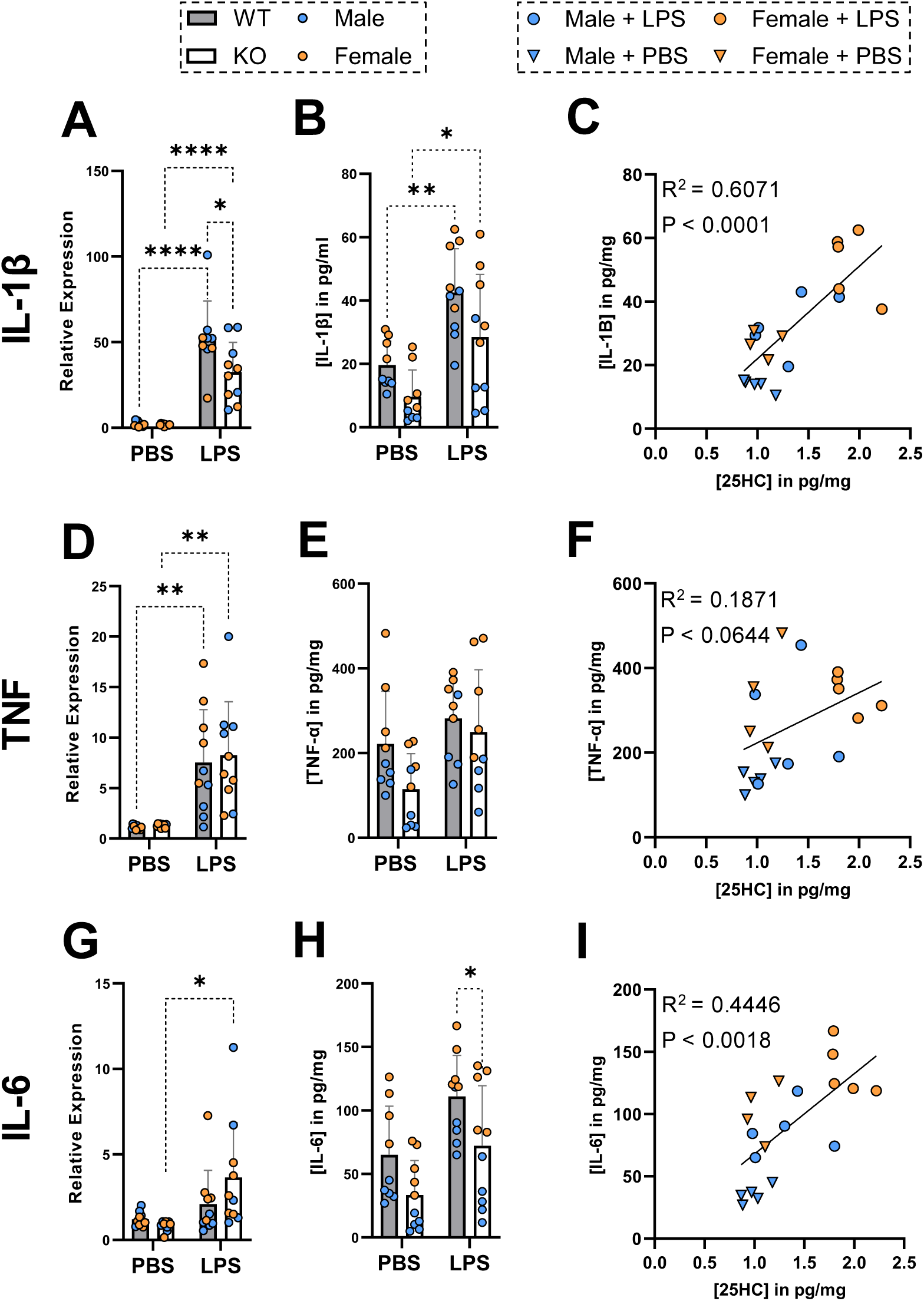
Levels of 25HC correlate with cytokine protein and gene expression. Relative expression of Il1b (A), Tnf (D) and Il6 (G) mRNA in hippocampus. Concentrations of IL-1β (B), TNF (E), and IL-6 (H) determined by ELISA and their respective correlations (C, F & I) with 25HC levels. 25HC concentrations are as shown in Figure 1C. Male mice are indicated in blue and female mice are in orange; PBS controls are inverted triangles and LPS-treated mice are circles – as shown at the top of the correlation plot. Data are presented as mean values ± SD. Statistical significance was defined using a two-way ANOVA with Tukey’s post-hoc test (*P < 0.05, **P < 0.01, & ****P < 0.0001). Significance and slope of the line was calculated with simple linear regression analysis and correlations were assessed by calculating the coefficient of determination (R2).

Measurement of IL-1β protein levels in brain extracts corroborated the mRNA findings (Fig 6B). Moreover, the brain levels of IL-1β showed a striking correlation with the levels of 25HC (from Fig 1C) (R^2^ = 0.607; p<0.0001) (Fig 6C). LPS-treated female mice that produced higher brain levels of 25HC also had higher brain levels of IL-1β relative to male mice.

Although expression of *Tnf* mRNA also significantly increased with LPS treatment (Fig 6D) in *WT* mice, this was not reflected by protein levels (Fig 6E). *Ch25h*-deficiency did not influence *Tnf* mRNA expression or TNFα protein levels in the brain. In addition, 25HC levels correlated weakly (R^2^ = 0.1871, p<0.0644) with TNFα protein levels (Figure 6F).

LPS treatment did not promote significant changes in *Il6* mRNA expression and IL6 protein levels (Figures 6G & H). However, significantly lower levels of IL6 protein were observed in *CKO* samples relative to *WT* after LPS treatment and 25HC levels correlated moderately with IL6 protein levels (Fig 6I, R^2^= 0.4446, p<0.0018).

These observations suggest that *Ch25h*-deficient mice present a reduced proinflammatory cytokine profile as predicted by the transcriptomic results.

### *Ch25h*-deficiency reduces hippocampal neutrophil infiltration following LPS treatment

Neutrophils comprise the bulk of circulating white blood cells and are the principal immune cells involved in the innate immune response to bacterial infection. The importance of neutrophil recruitment and TLR4 activation to an effective innate immune response to bacterial infection has been previously reported^31,32^. In the context of CNS bacterial infection, it has been demonstrated that neutrophil chemotaxis into the choroid plexus is TLR4-dependent^33^ and that LPS treatment using the same experimental paradigm used in our study, induces neutrophil infiltration into brain parenchyma^24^. Therefore, we examined the potential role of 25HC in the LPS-induced, TLR4-dependent CNS infiltration of neutrophils by measuring the presence of the neutrophil-specific marker lymphocyte antigen 6 complex locus G6D (LY6G) using qPCR and immunostaining in mouse hippocampus (Figure 7). In line with previous work^24^, *Ly6g* mRNA levels increased about 50-100-fold following LPS treatment in *WT* mouse brain. This increase was significantly reduced (p = 0.0067) in the *CKO* mice (Figure 7A). Intriguingly, females expressed higher levels of *Ly6g* mRNA than males upon LPS stimulation (Figure 7A).

**Figure 7:**
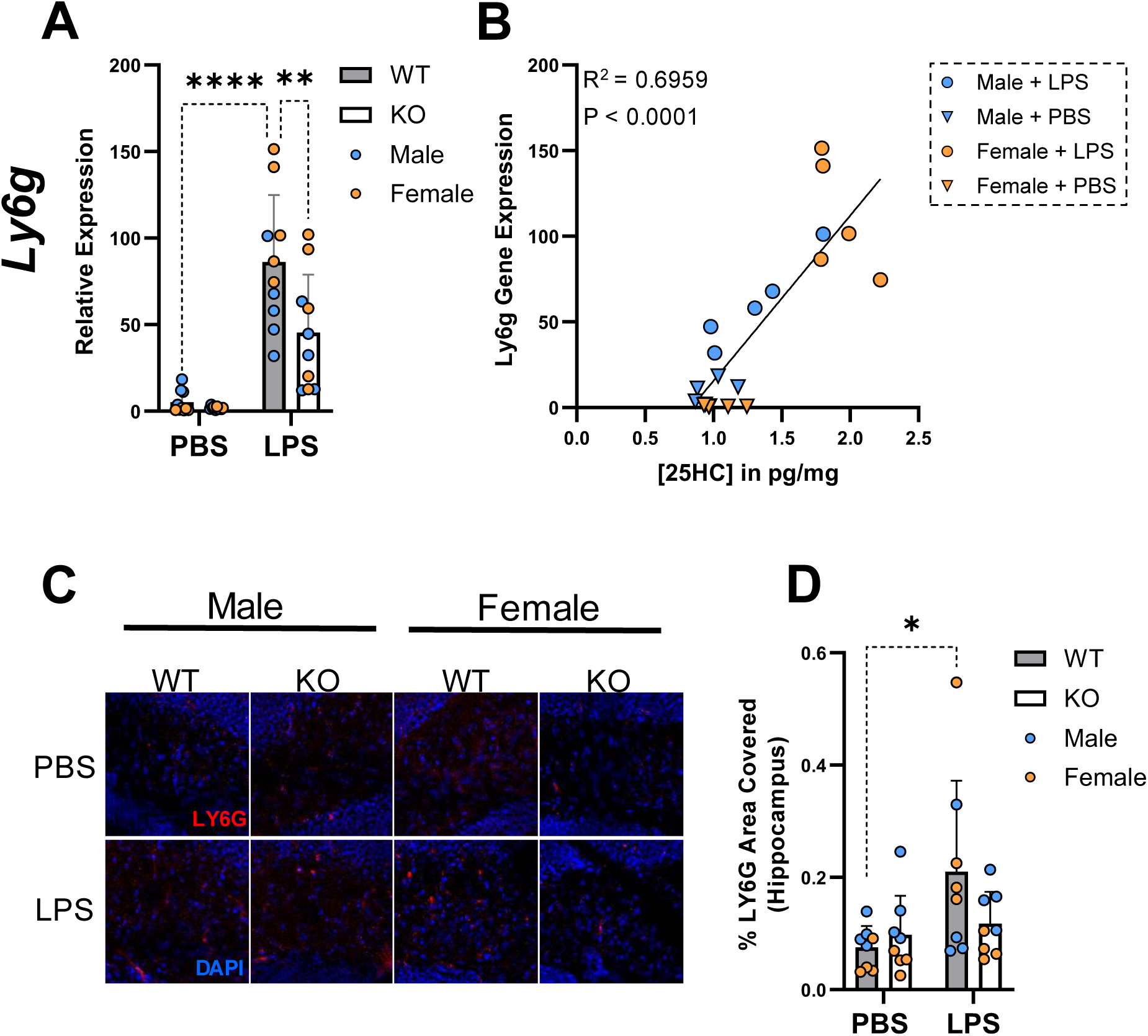
Levels of 25HC correlate with Ly6g gene expression and correspond to an increase in LY6G protein among WT mice. (A) Relative expression of Ly6g mRNA measured by qPCR in the hippocampus and (B) the correlation between Ly6g relative expression and 25HC levels. 25HC concentrations are as shown in Figure 1C. (C) Representative immunofluorescence images of mouse hippocampal sections stained for LY6G (red); nuclei were stained with DAPI (blue). (D) Percentage of area covered by LY6G in the hippocampus. Data are presented as mean values ± SD. Statistical significance was defined using a two-way ANOVA with Tukey’s post-hoc test (**P < 0.01 & ****P < 0.0001). Significance and slope of the line was calculated with simple linear regression analysis and correlations were assessed by calculating the coefficient of determination (R2).

Importantly, the brain concentrations of 25HC in *WT* mice (Fig 1C) showed a striking correlation (R^2^ = 0.696, p<0.0001) with relative expression of *Ly6g* mRNA. An even higher correlation (R^2^ = 0.933, p = 0.0075) was observed among the LPS-treated *WT* male mice, but a ceiling effect was observed among the LPS-treated female mice. To confirm if the increase in *Ly6g* mRNA corroborated with elevated levels of LY6G+ neutrophils in the brain parenchyma, we immunostained mouse brain tissue sections with an antibody specific to mouse LY6G (Figure 7C) and observed a significant increase (p < 0.05) in the percentage of area covered by LY6G immunoreactivity only in LPS-treated *WT* mice (Figure 7D). Differences in LY6G immunoreactivity between male and female mice were not significant with or without LPS treatment. Consistent with the lower levels of *Ly6g* mRNA expression observed in *CKO* mice (Figure 7A), LPS treatment in *CKO* mice did not result in a significant increase in LY6G immunoreactivity (Figure 7D).

## DISCUSSION

In this study, we use the “two-hit” model of intraperitoneal LPS administration^24^ to investigate the role of CH25H/25HC in LPS-induced neuroinflammation.

A growing body of research implicates neuroinflammation in the pathogenesis and progression of neurodegenerative disorders, such as AD^34,35^, with accumulating evidence suggesting roles for both innate and adaptive immunity as contributors^36–38^. There is also evidence that bacterial^39–41^ and viral^42,43^ infections as well as alterations in the microbiome^44^ may contribute to disease pathogenesis. Neuroinflammation often results when a peripheral infection spreads to the CNS. Activation of pattern-recognition receptors (PRRs) by pathogen-associated molecular patterns (PAMPs) occurs early in the brain’s innate immune response and to detect invading pathogens to initiate an inflammatory response. One class of PRRs, the Toll-like receptors (TLRs), are crucial for the innate immune response to bacterial infection. Toll-like receptor 4 (TLR4) canonically binds to the ubiquitously expressed gram-negative bacterial antigen, LPS, and is an important early trigger of inflammation in response to gram-negative bacterial infection. TLR4 inhibition or impairment has been shown to reduce brain inflammation in mouse models and influence the pathophysiology of neurodegenerative disease^45–47^. Recent work has found that periodontitis caused by bacterial infection of the gingiva is associated with an increased risk of AD^48^, suggesting that certain bacterial infections may increase the risk of developing AD. Given the prevalence of gram-negative bacteria in periodontitis resulting and its proximity to cerebral blood vasculature, the circulation of gram-negative bacteria (or LPS) near the blood-brain barrier is plausible. In the mouse model of neuroinflammation employed in our study, even peripheral LPS administration is sufficient to induce TLR4-mediated inflammation in brain parenchyma. As a cell-surface antigen that is ubiquitously expressed on gram-negative bacteria, LPS may play a role in mediating infection-related onset of neurodegenerative disorders like AD. Remarkably, even a single peripheral administration of LPS in wild-type mice has been reported to result in neuroinflammation that persists over an entire year^23^, suggesting similar mechanisms may prolong the inflammatory state in the CNS in humans.

Cholesterol 25-hydroxylase (CH25H) is markedly upregulated by LPS treatment and is one of the genes expressed by disease-associated microglia (DAM) that are chronically upregulated in various neurodegenerative disorders^10,11^. Prolonged expression of *Ch25h* and 25HC production may contribute to neuroinflammation and consequent neurodegeneration. In fact, we have recently observed that *Ch25h*/25HC deficiency markedly reduces the age-dependent neuroinflammation and neurodegeneration observed in the P301S tau transgenic mice (under review).

Reinforcing the findings from our immunochemical and qPCR assays, RNA-seq analysis of mouse hippocampal samples found that LPS significantly increased transcripts associated with immune-related biological pathways related to cytokine production, anti-viral immune response, immune cell activation, leukocyte infiltration, and inflammation more broadly.

While LPS treatment led to a significant increase in the microglial marker, IBA1 immunoreactivity in the hippocampus of both *WT* and *CKO* mice, *Ch25h* deficiency did not affect IBA1 immunoreactivity. However, in *Ch25h*-deficient mice there was a significant decrease in CD68 immunoreactivity, which is a marker of microglial phagolysosomes, suggesting that 25HC produced by *WT* microglia may limit the turnover of phagolysosomes. *Ch25h*-deficient mice also showed significantly decreased levels of the markers of microglial activation *Clec7a*, *Trem2* and *Axl*. Levels of 25HC in *WT* mice correlated strongly with IL1β and moderately with IL6 levels. Our previous studies with primary mouse microglia showed that not only did *Ch25h*-deficient microglia produce lower levels of IL1β but addition of 25HC resulted in increased IL1β production in response to LPS, suggesting that 25HC influences both expression and secretion of IL1β^12^. Intriguingly, recent work has found that *Ch25h*-deficient macrophages exhibited impaired mitochondria leading to leakage of mitochondrial DNA that in turn led to increased IL1β secretion via the AIM2 inflammasome^49^. The underlying reasons for the opposing effects of *Ch25h*-deficiency in microglia versus macrophages are unclear, but we speculate that other factors may influence myeloid cell physiology in a context- and cell- dependent manner.

Interestingly, brain levels of TNFα did not increase with LPS treatment, although prior research using a similar 1-hit LPS model had reported low but elevated TNF synthesis in mouse brain parenchyma using a more sensitive and precise cytometric bead assay^50^, suggesting that TNF was potentially upregulated at lower levels in our LPS-treated cohorts. Additionally, while cytokine secretion may have decreased over the prolonged LPS exposure over two days, expression of *Il1b, Il6, and Tnf* mRNAs remained upregulated by LPS treatment.

In addition to 25HC-mediated changes in inflammatory response by microglia, we also observed lower levels of astrocyte activation in LPS-treated *Ch25h*-deficient mice relative to wildtype mice. This underscores a potential role for 25HC in mediating microglia-astrocyte crosstalk. Many microglial factors including cytokines (IL1α, TNF) as well as other secreted factors have been shown to mediate astrocyte activation^51,52^. We recently showed that 25HC can dramatically modulate lipid metabolism in astrocytes via its effects in inhibiting SREBP-mediated gene expression and stimulating LXRs^14^. Thus, it is possible that changes in astrocyte lipid metabolism due to 25HC secreted by microglia might play a role in astrocyte activation and/or function.

In addition to the effects of 25HC on cytokine secretion, we have also provided evidence that 25HC strongly influences infiltration of neutrophils into the brain parenchyma. 25HC is metabolized by CYP7B1, a cytochrome P-450 enzyme, into 7α,25-dihydroxycholesterol (7α,25diHC)^53^ and 7α,25diHC is a potent ligand for the G-protein coupled receptor, GPR183/EBI2, which is expressed on leukocytes. 7α,25diHC activates GPR183/EBI2 to mediate chemotaxis of B-cells^4^ and infiltration of CD4+ T-cells into the brain parenchyma in a mouse model of experimental autoimmune encephalomyelitis (EAE)^54^. Comparing 2-day and 5-day serial administration of LPS every 24 hours, Thomson et al found dramatic increases in the presence of LY6G+ neutrophils that peaked at 2 days and had tapered off by 5 days of exposure to LPS while CD3+ T-cells showed the opposite trend increasing by 5 days of LPS^24^.

Intriguingly, while both male and female mice equally expressed markers of gliosis, the sexes diverged in cytokine production and neutrophil infiltration with females showing higher levels of both. Importantly, as the only known product of CH25H enzyme, levels of 25HC in brain were higher in females than males following LPS treatment. No sex differences were apparent in the low levels of 25HC that were independent of CH25H enzyme activity (i.e., 25HC levels that were observed in *ch25h*-/- mice). The observed sex differences in 25HC also correlated strongly with other neuroinflammatory markers and is commensurate with previous work observing relatively higher neuroinflammation in females^55^. Likewise, women have a well-established higher risk of neurodegenerative diseases such as AD that persists even after accounting for the greater life expectancy of women^56^. Whether 25HC contributes significantly to these sex differences in AD risk will require future investigation.

### Conclusions

Our findings demonstrate that 25HC is a sex-specific proinflammatory mediator of brain cytokine production and neutrophil infiltration in a mouse model of LPS-mediated neuroinflammation.

## MATERIALS AND METHODS

### Mouse model of LPS-induced neuroinflammation

Wild-type (C57BL/6J, #000664) and ch25h-/- (*CKO*, #016263) mice were purchased from The Jackson Laboratory and maintained as homozygotes. At 23-26 weeks of age, mice were intraperitoneally (i.p.) administered LPS (2 mg/kg per dose) or vehicle (PBS) twice within a 24-hours interval (Fig 1A). Animals were perfused 24 hours after the last injection. Left hemisphere was fixed in 4% paraformaldehyde, hippocampus and cortex were dissected from the right hemisphere and were frozen at −80°C until use. All animal procedures and experiments were performed under guidelines approved by the animal studies committee at Washington University School of Medicine.

### Immunofluorescence Staining

Left-hemisphere tissue sections from wild type and Ch25h knock out mice were sliced at 40 microns thickness using a cryostat (Thermo Fisher microm 430). Slices were then placed in a cryoprotectant (30% ethylene-glycol in PBS) and stored at −20°C until use in immunostaining. Two slices 960mm apart were taken for each animal (4 animals/treatment group). The sections were washed in Tris-buffered saline (TBS) 3 times for 5 minutes. Sections were then placed in a blocking solution made of 3% goat serum, 3% bovine serum albumin (BSA), and 0.25% Triton X-100 in TBS for 1 hour. Sections were then incubated with the primary antibody cocktail against CD68 (rat monoclonal; 1:250; catalog no. MCA1957; Bio-rad), GFAP (Chicken polyclonal; 1:1000; catalog no. Ab4674; Abcam), and IBA1 (Polyclonal rabbit; 1:500; catalog no. 019-19741; Waco) in TBS containing 1% goat serum and 1% BSA. The sections were left overnight at 4°C. The next day, tissue sections were washed three times with fresh TBS for 5 minutes each. The tissue sections were then placed in the secondary antibodies cocktail containing 1:1000 dilution each of DAPI, goat anti-rat-Alexa555 (Invitrogen; Catalog no. A21434), goat anti-chicken-Alexa 647 (Invitrogen; Catalog no. A21449), and goat anti-rabbit-Alexa488 (Invitrogen; Catalog no. A32731) in TBS. (Primary and secondary antibody cocktails were centrifuged at 10,000 rpm for 5 minutes to remove insoluble particulate matter prior to use. The stained sections were then washed three times with TBS for 7 minutes each. The slices were mounted in low light in mounted using FluoroShield mounting media (Sigma-Aldrich; Product no. F6182).

### Imaging

Images were retrieved using a Nikon spinning disk confocal microscope. Using Fiji, a region of interest (ROI) around the hippocampus was drawn. Background subtraction was conducted using a 50-pixel rolling ball. Using batch processing, same threshold settings were applied to all the images of each individual stain and area fraction was measured.

### Tissue Extract preparation

From each brain, ∼10-15mg of mouse hippocampal and cortical tissue were sliced on a dry ice chilled glass plate, placed in 1.5 mL Eppendorf tubes. Frozen cortical tissue was homogenized with a micro pestle and electric rotor in 200 μL of mammalian protein extraction reagent (M-PER) (Thermo Scientific, Cat. #: 78501) containing ProBlock Gold Extra Strength Protease Inhibitor Cocktail Kit (GoldBio, Cat. #: GB-116-2). The cortical lysate was then aliquoted into a 96 well plate and stored at −80°C.

### Enzyme-Linked Immunosorbent Assay (ELISA)

ELISA kits from R&D Systems were used to quantify various cytokines, including IL-1B (Cat #: DY401-05), TNF-a (Cat #: DY410-05), and IL-6 (Cat #: DY40605) in the posterior cortical lysates of the *WT* and *CKO* mice. The assays were performed as instructed in the manufacturer’s protocols, except ELISA steps were performed in half-well rather than full-well plates (Corning Cat#: 3690) with half of the volume in each well (50 μL instead of 100 μL).

### RNA Extraction

Frozen hippocampal tissue was homogenized with a micropestle and electric rotor in RNA Lysis Buffer from Zymo Research (Cat#: R2060-1-100) before immediately proceeding with RNA extraction using the Zymo Research Quick-RNA MiniPrep Plus kit (Cat. #: R1058) according to the manufacturer’s instructions. After eluting the RNA into nuclease-free water, the RNA concentrations were measured using a NanoDrop spectrophotometer.

### Reverse Transcription cDNA Synthesis & Quantitative Polymerase Chain Reaction (qPCR)

The High-Capacity cDNA Reverse Transcription with RNase Inhibitor Kit (Applied Biosystems, Cat. #: 4474966) was used to reverse transcribe 100-200ng of RNA in accordance with the manufacturer’s instructions. For the qPCR reaction mix, PrimeTime probe-based qPCR assays (see Supplementary Table V) and PrimeTime Gene Expression Master mix (IDT, Cat# 1055771) were obtained from IDT (Integrated DNA Technologies, Inc). qPCR reactions were run using the “Fast” mode on a QuantStudio™ 3 Real-Time PCR Instrument (Applied Biosystems by ThermoFisher, A28131). Probes used for various genes are listed in Table X. Data were normalized against actin (Actb) and the 2^−ΔΔCt^ method was used to calculate relative gene expression value (Relative Quantification, RQ).

### RNA Sequencing and Analysis

Samples were prepared according to library kit manufacturer’s protocol, indexed, pooled, and sequenced on an Illumina NovaSeq 6000. Basecalls and demultiplexing were performed with Illumina’s bcl2fastq2 software. RNA-seq reads were then aligned and quantitated to the Ensembl release 101 primary assembly with an Illumina DRAGEN Bio-IT on-premise server running version 3.9.3-8 software.

All gene counts were then imported into the R/Bioconductor package EdgeR^57^ and TMM normalization size factors were calculated to adjust for samples for differences in library size. Ribosomal genes and genes not expressed greater than one count-per-million in the smallest group size minus one samples were excluded from further analysis. The TMM size factors and the matrix of counts were then imported into the R/Bioconductor package Limma^58^. Weighted likelihoods based on the observed mean-variance relationship of every gene and sample were then calculated for all samples and the count matrix was transformed to moderated log 2 counts-per-million with Limma’s voomWithQualityWeights^59^. The performance of all genes was assessed with plots of the residual standard deviation of every gene to their average log-count with a robustly fitted trend line of the residuals. Differential expression analysis was then performed to analyze for differences between conditions and the results were filtered for only those genes with Benjamini-Hochberg false-discovery rate adjusted p-values less than or equal to 0.05. For each contrast extracted with Limma, biological enrichment was carried out with the web tool “Metascape” to identify significantly enriched Gene Ontology (GO) terms {Zhou et al., 2019, #180370}. Pathway maps were generated using Cytoscape ^61^.

### Liquid Chromatography-Mass Spectrometry (LC/MS)

Brain samples from 40 mice were profiled for oxysterols (3β, 5α, 6β-trihydroxycholestane (Triol), and 7-ketocholesterol (7-KC), 4β-hydroxycholesterol (4β-HOC), 7αhydroxycholesterol (7α-HOC), 7β-hydroxycholesterol (7β-HOC), 24-hydroxycholesterol (24-HOC), 25-hydroxycholesterol (25-HOC), and 27-hydroxycholesterol (27-HOC)). The brain samples were homogenized in water (20 mL/g tissue) using Omni Bead Ruptor 24 (Omni International, Inc.).

The oxysterols in 50 µL of homogenate were extracted with liquid-liquid extraction after addition of Triol-d7, 7-KC-d7, 4β-HOC-d7, 7α-HOC-d7, 7βHOC–d7, 24-HOC–d7, 25-HOC-d6, 27-HOC–d5 as the internal standards. The oxysterols and their internal standards were derivatized with nicotinic acid to increase their mass spectrometric sensitivities. Quality control (QC) samples were prepared by pooling a portion of study samples, and injected every 5 samples to monitor instrument performance. The sample analysis was performed with a Shimadzu 20AD HPLC system coupled to a 4000QTRAP mass spectrometer operated in positive multiple reaction monitoring mode. Data processing was conducted with Analyst 1.6.3. The relative quantification data were reported as peak area ratios of analytes to their internal standards.

## LEGENDS FOR SUPPLEMENTARY FIGURES

**Figure S1:**
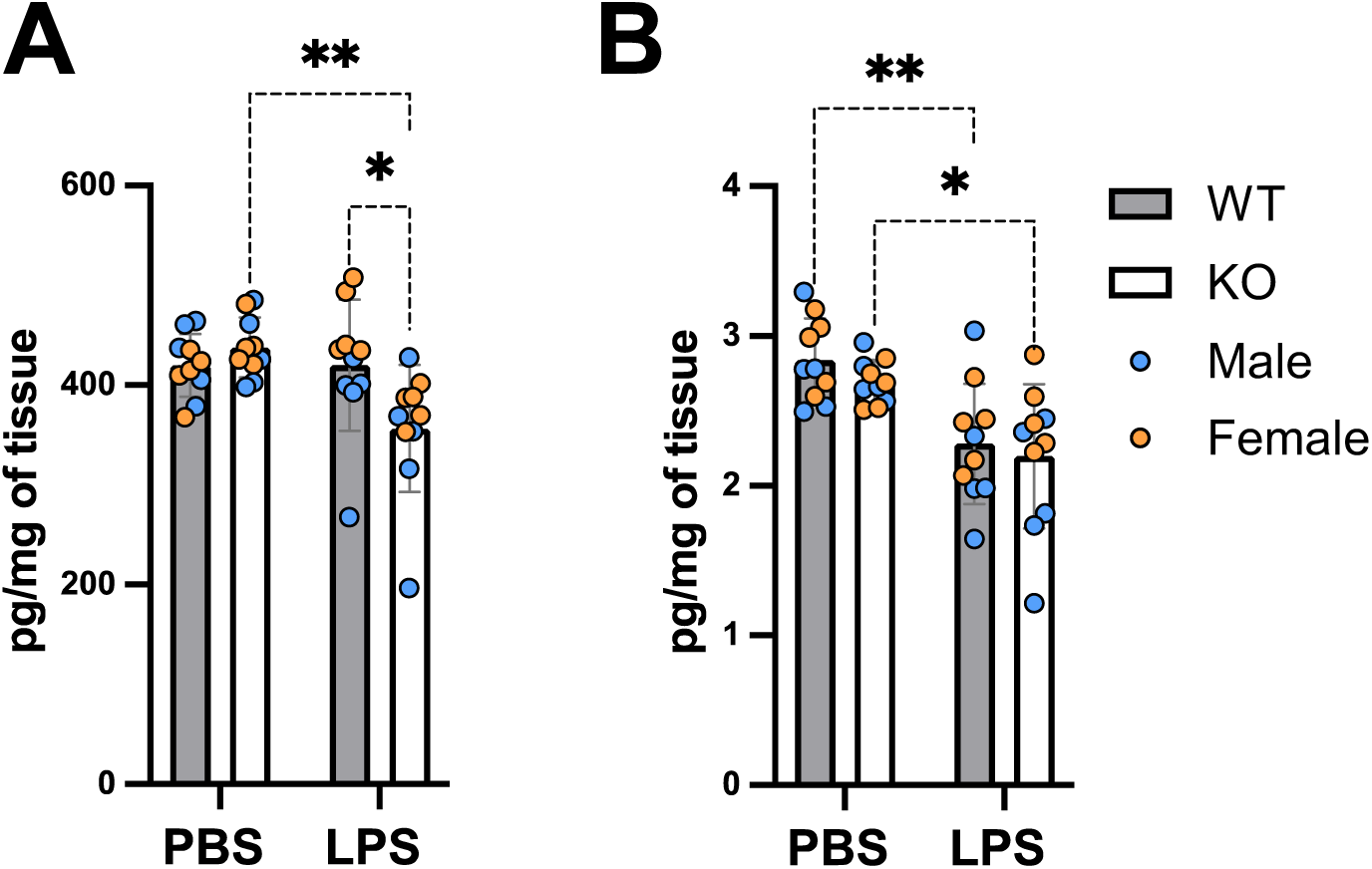
Quantification of the two structural isomers of 25HC. Quantitation of 24-(S)-HC (A) and 27-HC (B) in PBS- and LPS-injected wildtype (WT) and Ch25h-deficient (KO) mouse brains. Male mice are indicated in blue and female mice are in orange. Data are presented as mean values ± SD. Statistical significance was defined using a two-way ANOVA with Tukey’s post-hoc test (*P < 0.05 & **P < 0.01).

**Figure S2:**
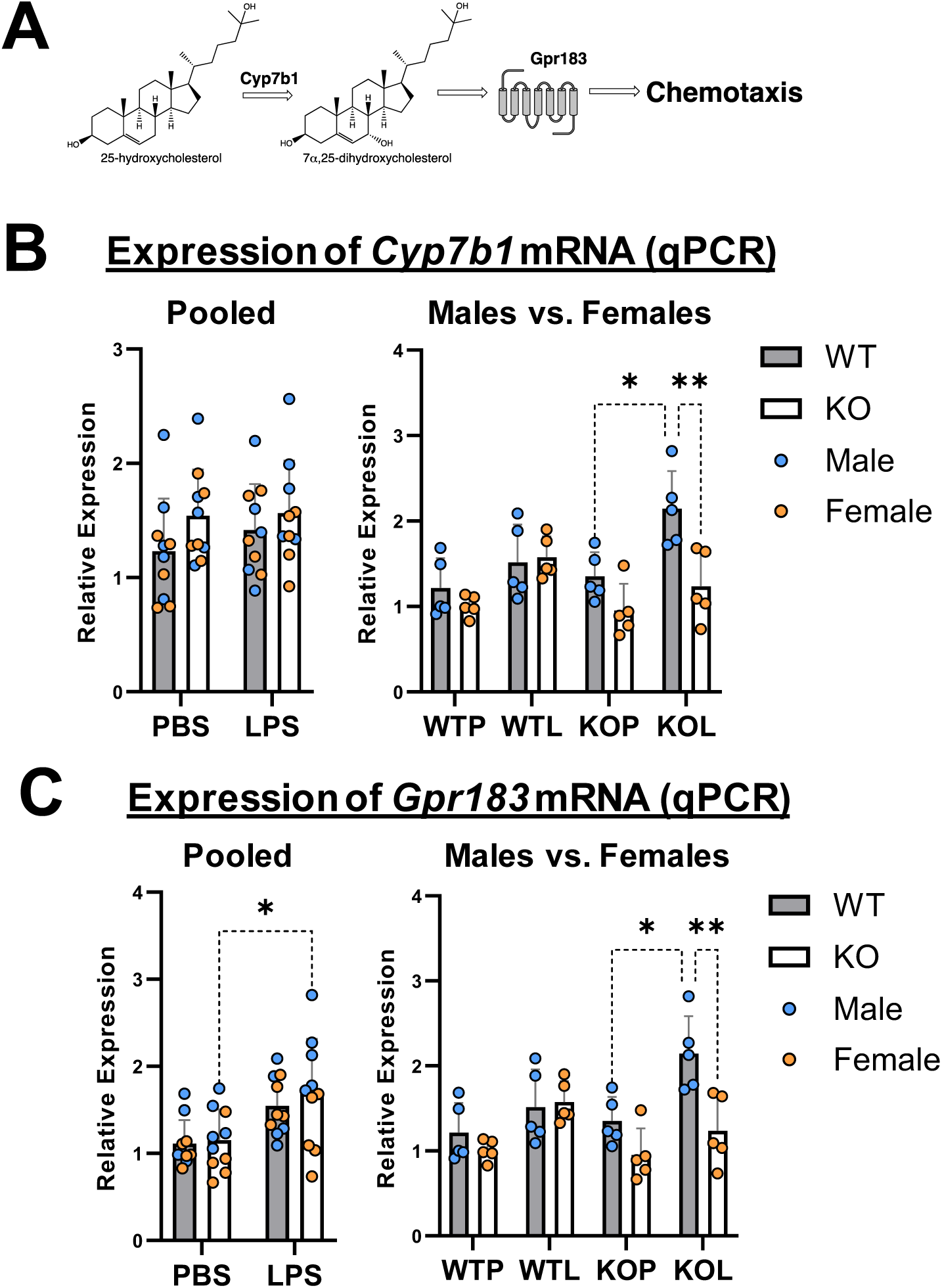
Expression of Cyp7b1 and Gpr183. (A) Schematic of 25HC pathway leading to chemoattraction of GPR183 expressing leukocytes. (B) Relative expression of Cyp7b1 mRNA measured by qPCR - pooled (left) and sex-segregated (right). (C) Relative expression of Gpr183 mRNA measured by qPCR - pooled (left) and sex-segregated (right). Data are presented as mean values ± SD. Statistical significance was defined using a two-way ANOVA with Tukey’s post-hoc test (*P < 0.05, **P < 0.01).

**Figure S3:**
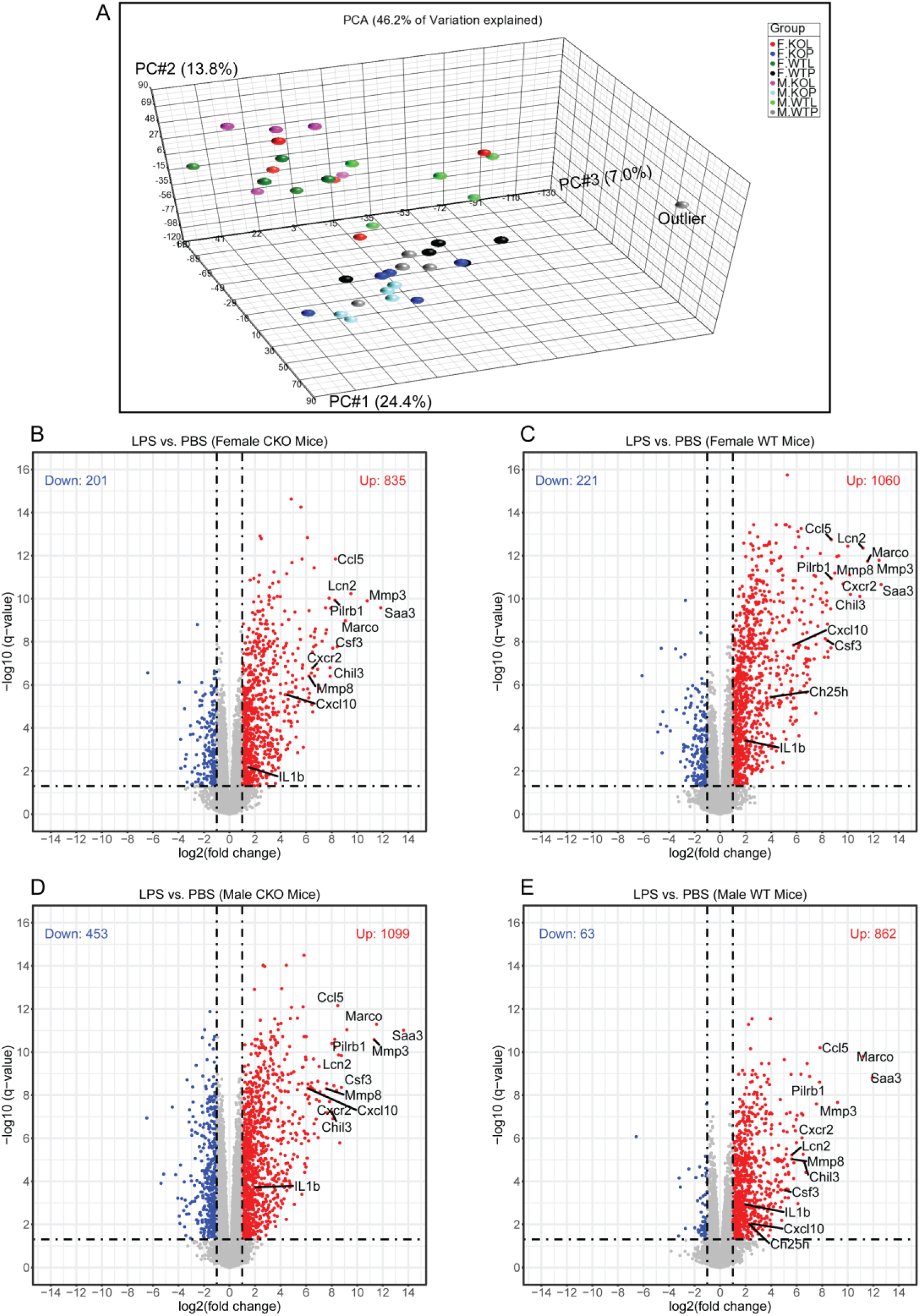
Transcriptomic data. (A) Principal component analysis (PCA) of RNA-seq measurements from 40 experimental mice in female and male treated by LPS or PBS. One sample was an outlier and was excluded from further analysis. (B-E) Volcano plots of differentially expressed genes (DEGs) in these animals. DEGs were defined as those with fold- change >= 2 at adjusted p-value (q-value) < 0.05.

**Figure S4:**
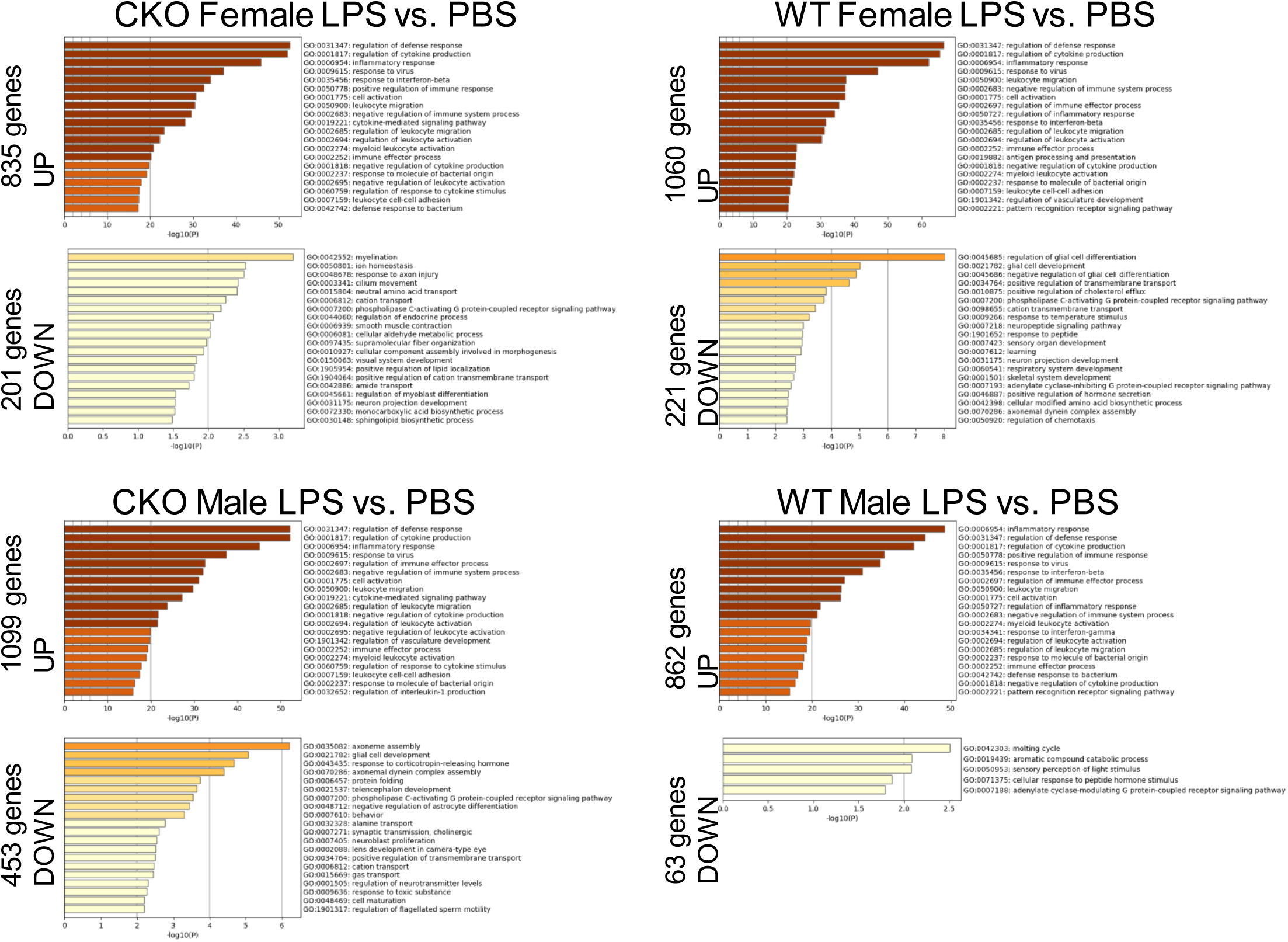
Bar graph of enriched terms across input gene lists, colored by p-values. P-values are also shown on the X-axis.

## SUPPLEMENTARY TABLES

**Supplementary Table I:** Size of gene-lists influenced by gender and genotype

**Supplementary Table II:** Differential gene expression between females and males

**Supplementary Table III:** Gene ontology analysis of differentially expressed genes

**Supplementary Table IV:** Comparisons of the downregulated genes and pathways in response to LPS treatment

**Supplementary Table V:** Sequences of primers used for qPCR analysis.

## ABBREVIATIONS

LPS: Lipopolysaccharide
AD: Alzheimer’s disease
TLR4: Toll-like receptor-4
CH25H: Cholesterol-25-hydroxylase
25HC: 25-hydroxycholesterol
7α,25diHC: 7α,25-dihydroxycholesterol
DAM: Disease-associated microglia,

## DECLARATIONS

### Ethics approval and consent to participate

Research using mouse as a model system of neuroinflammation was approved by the Institutional Animal Care and Use Committee (IACUC) of the Washington University School of Medicine.

### Consent for publication

Not applicable

### Availability of data and materials

All data generated or analysed during this study are included in this published article and its supplementary information files.

### Authors’ contributions

AGC: conceived and designed the research studies; JR, DTR, AGC: performed experiments and acquired data; JR, DTR, JY, AGC: analyzed the data; JR, AGC: wrote the paper; JR, DTR, SMP, AGC: edited the manuscript.

## Acknowledgements

We acknowledge technical assistance from Kaylee Stillwell and from Washington University core facilities. We also acknowledge discussions with members of Sage Therapeutics, Cambridge, MA.

